# Targeted delivery of acid alpha-glucosidase corrects skeletal muscle phenotypes in Pompe disease mice

**DOI:** 10.1101/2020.04.22.051672

**Authors:** Andrew D. Baik, Philip T. Calafati, Nina A. Aaron, Antonia Mehra, Sven Moller-Tank, Lawrence Miloscio, Lili Wang, Maria Praggastis, Matthew S. Birnbaum, Cheryl Pan, Susannah Brydges, Alejandro Mujica, Peter Barbounis, Nicholas W. Gale, Ning Li, Christos A. Kyratsous, Christopher J. Schoenherr, Andrew J. Murphy, Aris N. Economides, Katherine D. Cygnar

## Abstract

Lysosomal diseases are a class of genetic disorders predominantly caused by loss of lysosomal hydrolases, leading to lysosomal and cellular dysfunction. Enzyme Replacement Therapy (ERT), where recombinant enzyme is given intravenously, internalized by cells, and trafficked to the lysosome, has been applied to treat several lysosomal diseases. However, current ERT regimens do not correct disease phenotypes in all affected organs because the biodistribution of enzyme uptake does not match that of the affected cells and tissues that require the enzyme. We present here targeted ERT, an approach that utilizes antibody-enzyme fusion proteins to target the enzyme to specific tissues. The antibody moiety recognizes transmembrane proteins involved in lysosomal trafficking and that are also preferentially expressed in those cells most affected in disease. Using Pompe disease (PD) as an example, we show that targeted ERT is superior to ERT in treating the skeletal muscle phenotypes of PD mice.

## Introduction

Pompe disease (PD), or Glycogen Storage Disease type II, is a monogenic, lysosomal disease caused by a deficiency in the activity of the enzyme lysosomal acid alpha-glucosidase (GAA). GAA deficiency results in an accumulation of its substrate, glycogen, in the lysosomes of tissues including skeletal and cardiac muscle. This aberrant accumulation of glycogen in myofibers results in progressive damage of muscle tissue, with symptoms that can include cardiomegaly, mild to profound muscle weakness, and ultimately death to due cardiac or respiratory failure^1^.

As is the case for several other lysosomal diseases, PD is currently treated by enzyme replacement therapy (ERT). For PD, recombinant human GAA (rGAA) is delivered by intravenous infusion into patients every other week. While ERT has been very successful in treating the cardiac manifestations of PD, skeletal muscle and the central nervous system (CNS) remain minimally treated by ERT ^2–4^.

Inadequate delivery of rGAA to skeletal muscle has been suggested as the main reason ERT fails to treat this organ system ^5^. The primary mechanism by which rGAA reaches lysosomes is through uptake by the cation-independent mannose 6-phosphate receptor (CI-MPR), which binds mannose 6-phosphate on rGAA. However, CI-MPR expression in skeletal muscle is very low, and rGAA is poorly mannose 6-phosphorylated ^6^. In addition, CI-MPR may be misdirected into autophagosomes in affected cells, rather than lysosomes^7^, while a large amount of the drug is also taken up by liver (an organ that does not have primary pathology in PD). To overcome these problems, rGAA is given at a high and frequent dosing (20-40 mg/kg once every two weeks); nonetheless, difficulties in treating skeletal muscle persist. Strategies to modify rGAA to increase delivery to muscle by manipulating the affinity of GAA to CI-MPR, such as increasing the mannose 6-phosphate content of GAA^8^ or using the IGF2 binding sites on CI-MPR9 have been explored in the clinic. Still, these methods fail to address the underlying inherent limitations of using CI-MPR as the mechanism for uptake*, i.e.* CI-MPR’s low level of muscle expression and its largely endosomal rather than surface localization in skeletal muscle, which combined with high expression in the liver, result in very poor uptake by that tissue^10,11^.

The poor uptake by skeletal muscle, is further exacerbated by significant immune responses to rGAA, including neutralizing antibodies, which further reduce the effectiveness of ERT in a subset of patients^12,13^. Recently, liver-restricted expression of GAA has been shown to tolerize GAA knockout (KO) mice to GAA and mitigate the antibody immune response to GAA in the serum ^14^. However, while the immune response was dampened, correction of skeletal muscle glycogen accumulation was only achieved at very high doses of AAV infection^15,16^, likely a reflection of the continued reliance on CI-MPR for uptake by skeletal muscle. It was also unclear what proportion of hepatically secreted GAA is mannose 6-phosphorylated, as overexpression of GAA in other cell systems has been shown to adversely affect mannose 6-phosphate content of GAA^17,18^.

Therefore, the current state of the art is one of only incremental progress in spite of continued attempts at improving ERT. We propose that further improvements in ERT require new technological approaches that directly, and preferably concurrently, address the two main limitations of current ERT – i.e. effective delivery to affected tissues and immunogenicity. Using PD as an example, we present here targeted ERT, a new technology which addresses the delivery and biodistribution limitations of ERT by employing antibody-enzyme fusion proteins as the drug and testing them in PD mice. In targeted ERT, the antibody moiety targets the enzyme, GAA, to the affected tissues, thereby achieving superior clearance of the substrate – glycogen – compared to non-targeted GAA. To address the immunogenicity aspect of ERT, we adapt targeted ERT to AAV-mediated gene delivery to the liver. This not only renders it less immunogenic, but also provides long-term efficacy while also alleviating the need for repeated dosing and prophylaxis from anaphylaxis in mice and positions targeted ERT as a modality that can be adopted in emerging gene therapy-based treatments for LDs.

## Results

### Antibody:GAA fusion proteins display equivalent enzymatic activity to GAA

In order to address the delivery limitations associated with GAA ERT – *i.e.* loss of drug to the liver and poor delivery to skeletal muscle – we chose to bypass CI-MPR-mediated delivery of GAA and explore antibody-enzyme fusion proteins as a means to target the enzyme to the affected tissues. These fusion proteins have to display the following properties:

1. *Retain enzymatic function*.
2. *Bind to effector proteins that traffic to the lysosome*, most likely transmembrane or cell surface proteins that are also preferentially expressed in the tissues most affected in disease, while displaying minimal or no expression in non-affected tissues and acting independently of CI-MPR.

In order to identify proteins with the desired properties specifically for GAA ERT, we screened an internal database of gene expression profiles and defined a set of transmembrane and cell surface proteins that are expressed in skeletal muscles, and which are minimally expressed in the liver. We then looked among them for those with monoclonal antibodies readily available. Two of these proteins, CD63 (Figure S1A, B) and ITGA7 ^17^, met all three criteria. Furthermore, CD63 was already known to traffic between the cell surface and lysosomes ^18,19^, suggesting that it could be used as a benchmark to compare other potential effectors.

Starting with CD63, we engineered antibody:GAA fusions where the C-terminus of full-length IgG4 antibody was fused to the N-terminus of amino acids 70-952 of GAA using a glycine-serine linker (Figure S1C). The resulting antibody:GAA fusion, α-hCD63:GAA, was produced in CHO cells and purified by protein A chromatography. No cleavage products were seen in the supernatant, indicating that the α-hCD63:GAA fusion remained intact (Figure S1D). The enzymatic activity of purified α-hCD63:GAA fusions was found to be comparable to rGAA alone using the 4-methylumbelliferyl-alpha-D-glucoside fluorogenic substrate for the enzyme *in vitro* (Figure S1E).

### Internalization of antibody:GAA fusions is an antibody-dependent, CI-MPR-independent process

To verify the antibody-mediated properties the antibody:GAA fusions, the α-hCD63:GAA fusion was incubated with HEK293 cells overnight to allow for intracellular uptake. Dose-dependent cellular uptake was observed, while a non-binding antibody:GAA control showed no uptake in these cells. Furthermore, α-hCD63:GAA did not internalize in HEK293 cells that are deficient for hCD63. These results demonstrated that α-hCD63:GAA internalized specifically through the effector (CD63) recognized by the antibody domain of the fusion protein (Figure 1A).

**Figure 1.**
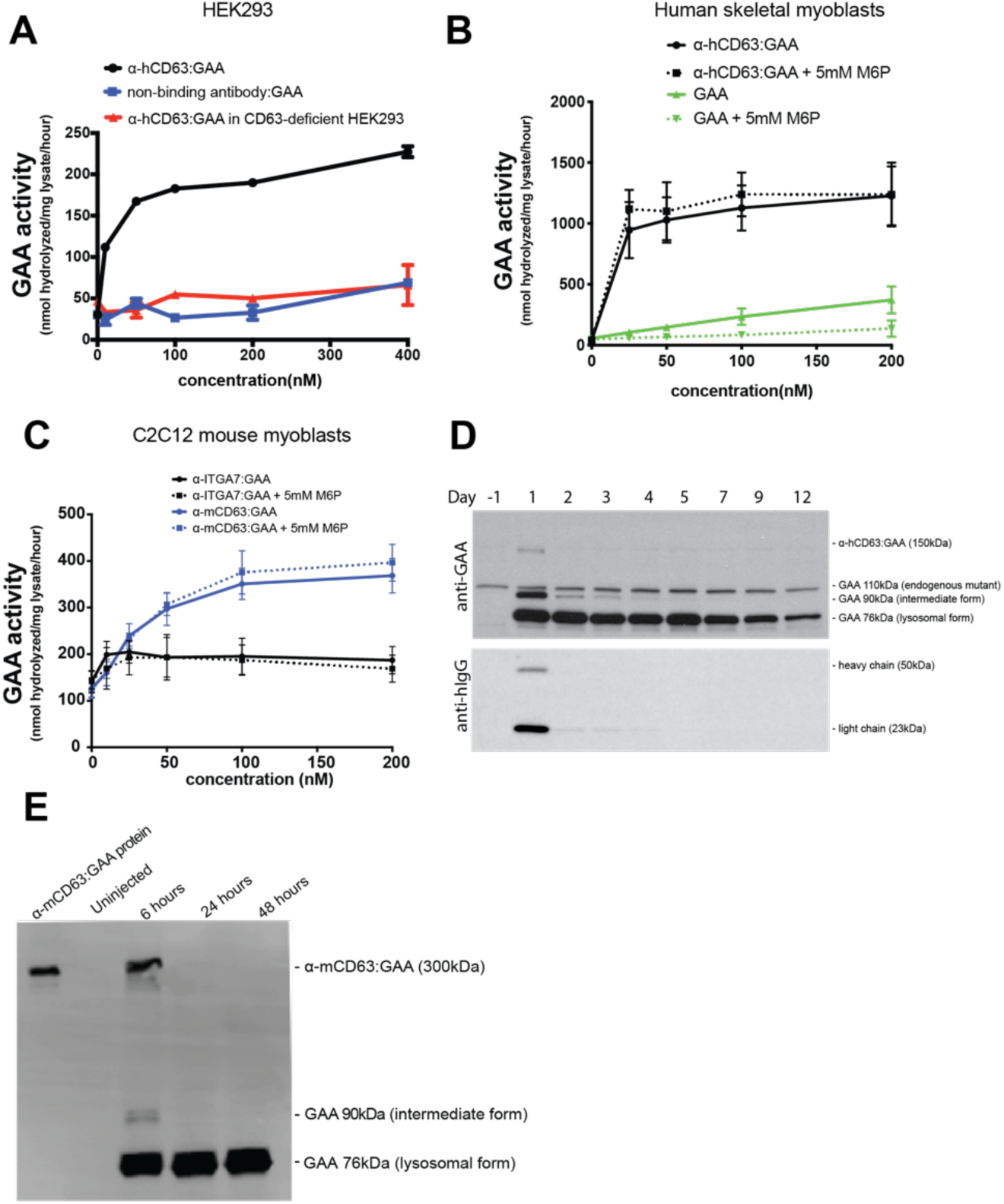
Antibody:GAA proteins are endocytosed by an antibody-mediated, effector-dependent and CI-MPR-independent mechanism and are processed post-internalization to the mature, lysosomal GAA form. (A) α-hCD63:GAA is internalized via CD63. GAA activity was detected in HEK293 cell lysate after overnight incubation of cells with α-hCD63:GAA. Eliminating the interaction between the antibody and CD63, either by using an IgG4 isotype control fused to GAA (IgG4:GAA) or CD63-deficient HEK293 cells reduced the GAA activity in the lysate. (B) Saturation of CI-MPR by addition of free M6P blocks uptake of GAA but does not block uptake of α-hCD63:GAA in human skeletal myoblasts or (C) block uptake of α-mCD63:GAA in C2C12 mouse myoblasts. (D) α-hCD63:GAA is processed into the mature GAA form and persists for at least 12 days in Pompe patient fibroblasts (probed with anti-GAA, upper panel). The antibody portion, which was cleaved off, is rapidly degraded (probed with anti-hIgG, lower panel). Note that these Pompe patient-derived fibroblasts lack the lysosomal forms but retain a 110kDa mutant endogenous form of GAA, as seen in lane “Day −1”. (E) α-mCD63:GAA is processed to the mature, lysosomal form *in vivo* (liver lysates from a PD mouse probed with α-GAA 6, 24, and 48 hours after injection). Error bars mean are +/− SD.

A glycan analysis (Figure S2) of α-hCD63:GAA showed that M6P was not present in any of the N-linked glycosylation sites present on this fusion protein. To verify that CI-MPR is not involved in the uptake in a relevant cell type, α-hCD63:GAA was assayed for internalization in human primary myoblasts (Figure 1B). Uptake saturated at ~50nM and was not inhibited by the presence of 5mM mannose 6-phosphate (M6P), a competitive inhibitor for CI-MPR binding. The ability of 5mM M6P to inhibit CI-MPR mediated uptake of GAA was verified in the same cell line.

To explore whether the properties observed with α-hCD63:GAA are shared when other effectors are utilized, we tested an α-ITGA7:GAA fusion protein. This targets mouse integrin alpha-7, a surface protein preferentially expressed in muscle tissue ^20^. α-ITGA7:GAA was incubated with C2C12 mouse myoblasts overnight and compared to α-mCD63:GAA. Both constructs were internalized in C2C12s independently of M6P, as 5mM M6P did not decrease uptake (Figure 1C). However, α-mCD63:GAA exhibited higher internalization than α-ITGA7:GAA.

### Antibody:GAA fusions undergo normal intracellular processing of GAA

To determine if the precursor GAA on the α-hCD63:GAA fusion retains the intracellular proteolytic processing events to form mature lysosomal forms of GAA ^21^, we performed a pulse-chase experiment of α-hCD63:GAA on a Pompe patient fibroblast line that lacks the lysosomal GAA forms. Within 24 hours of internalization, accumulation of the 76kDa lysosomal form of GAA was observed (Figure 1D). The half-life of the lysosomal GAA matched the long intracellular half-life of other rGAA preparations ^9^, while the antibody portion of the fusion quickly degraded over 3 days. Cathepsin inhibitors known to abolish the processing of GAA to the lysosomal form prevented full processing of the antibody:GAA to the lysosomal 76 kDa form (Figure S3) ^22^, further indicating that the α-hCD63:GAA fusion protein is converted in the lysosome into mature GAA. Importantly, the processing of antibody:GAA fusions to the lysosomal GAA form was further demonstrated *in vivo* (Figure 1E). In a mouse model of PD, *Gaa*^*6neo/6neo*^ ^23^, a single intravenous dose of 50mg/kg α-mCD63:GAA protein results in accumulation of the lysosomal form of GAA within 6 hours of injection (the earliest time point tested) and persists for at least 48 hours post-injection (*i.e.* at the latest time point tested).

### Targeted GAA has superior in vivo efficacy versus GAA alone

In the absence of access to the actual rGAA utilized in the clinic, we opted to test the efficacy of targeted GAA versus GAA using hydrodynamic delivery (HDD), a method that introduces plasmid DNA into mouse hepatocytes *in vivo* ^24^. We administered expression vectors encoding either α-mCD63:GAA, an isotype control antibody:GAA, or GAA to 2 to 3 month old *Gaa*^*6neo/6neo*^ mice and assessed GAA levels in serum and clearance of muscle glycogen 3 weeks post-HDD. Serum levels of α-mCD63:GAA, isotype control antibody:GAA, or GAA protein were roughly equal in the corresponding groups and detectable up to 10 days (Figure S4A). In the heart, both α-mCD63:GAA and GAA appeared to be equally efficacious as they reduced glycogen to wild-type levels. In contrast, in all skeletal muscles assayed, α-mCD63:GAA removed more glycogen than GAA, demonstrating that delivery to muscles was enhanced by the antibody moiety of α-mCD63:GAA (Figure S4B). As a negative control, an isotype control antibody:GAA with no known binding to any proteins in the mouse proteome did not clear glycogen. This showed that antibody-mediated targeting to CD63, rather than some other property of α-mCD63:GAA, was responsible for the observed clearance of glycogen.

In addition to CD63, we tested ITGA7 as an additional effector, using an α-ITGA7:GAA fusion protein, and obtained similar results (Figure S4C). This indicates that this approach is not limited to CD63 and that other transmembrane proteins can be used as effectors. The differences in *in vitro* and *in vivo* uptake/efficacy data of the α-ITGA7:GAA may be due to different kinetics and expression levels of *Itga7* in mouse cell lines versus mouse tissues.

### scFv:GAA clears glycogen similarly to antibody:GAA

In order to address the size constraints of AAV packaging for gene therapy and facilitate clinical translation of targeted GAA either as protein or by gene therapy, we explored GAA fusions with human α-hCD63 antibodies in two different formats, full length antibody or an scFv (Figure S1C). Given that these fusions recognize human, but not mouse, CD63, we engineered another PD mouse model, *Gaa*^*−/−*^;*Cd63*^*hu/hu*^, where *Gaa* was replaced with an ORF encoding LacZ and the protein-coding region of the *Cd63* locus has been replaced with its human counterpart (Figure S5). *Gaa*^*−/−*^;*Cd63*^*hu/hu*^ mice displayed equivalent levels of glycogen storage as *Gaa*6neo/6neo mice (Figure S6). With this humanized *Cd63* model of PD in place, we compared α-hCD63 in an scFv:GAA format (hereon referred to as α-hCD63_SC_:GAA) with α-hCD63:GAA to clear glycogen, using HDD to introduce the respective constructs. α-hCD63_SC_:GAA removed glycogen to a similar extent as the full-length antibody version (Figure S4C). This result cleared the way for more detailed comparisons of α-hCD63_SC_:GAA with GAA using AAV-mediated gene therapy, as full length antibody:GAA fusion transgenes are too large to package into AAV.

### *AAV α*-hCD63_SC_*:GAA is expressed more efficiently than AAV GAA from the liver*

In order to achieve long-term expression and sustained delivery of GAA and antibody:GAA proteins, we utilized AAV-mediated liver depot gene therapy in *Gaa*^*−/−*^;*Cd63*^*hu/hu*^ mice. AAV2/8 viruses were packaged with cDNA encoding α-hCD63_SC_:GAA or GAA driven by a liver-specific promoter, AAV α-hCD63_SC_:GAA and AAV GAA, respectively. The AAVs were injected by tail-vein into 2 to 3 month old *Gaa*^*−/−*^;*Cd63*^*hu/hu*^ mice at 1e10 or 1e11 vg/mouse. Serum levels of GAA proteins were higher in the AAV α-hCD63_SC_:GAA groups than in the AAV GAA-treated groups at equivalent doses (Figure 2A). RNA expression levels were measured in lysates of heart, quadriceps, and liver 3 months after infection, and expression was predominant in the liver for all constructs in a viral dose-dependent fashion (Figure S7). Interestingly, the ratio of secreted protein to RNA expression was much higher in the α-hCD63_SC_:GAA groups (Figure 2B). We explored secretion differences of the two constructs *in vitro* by transfecting the Huh-7 hepatocyte cell line with α-hCD63_SC_:GAA or GAA. A higher ratio of secreted to intracellular translated protein was observed in the α-hCD63_SC_:GAA transfected cells than in the GAA transfected cells (Figure S8).

**Figure 2.**
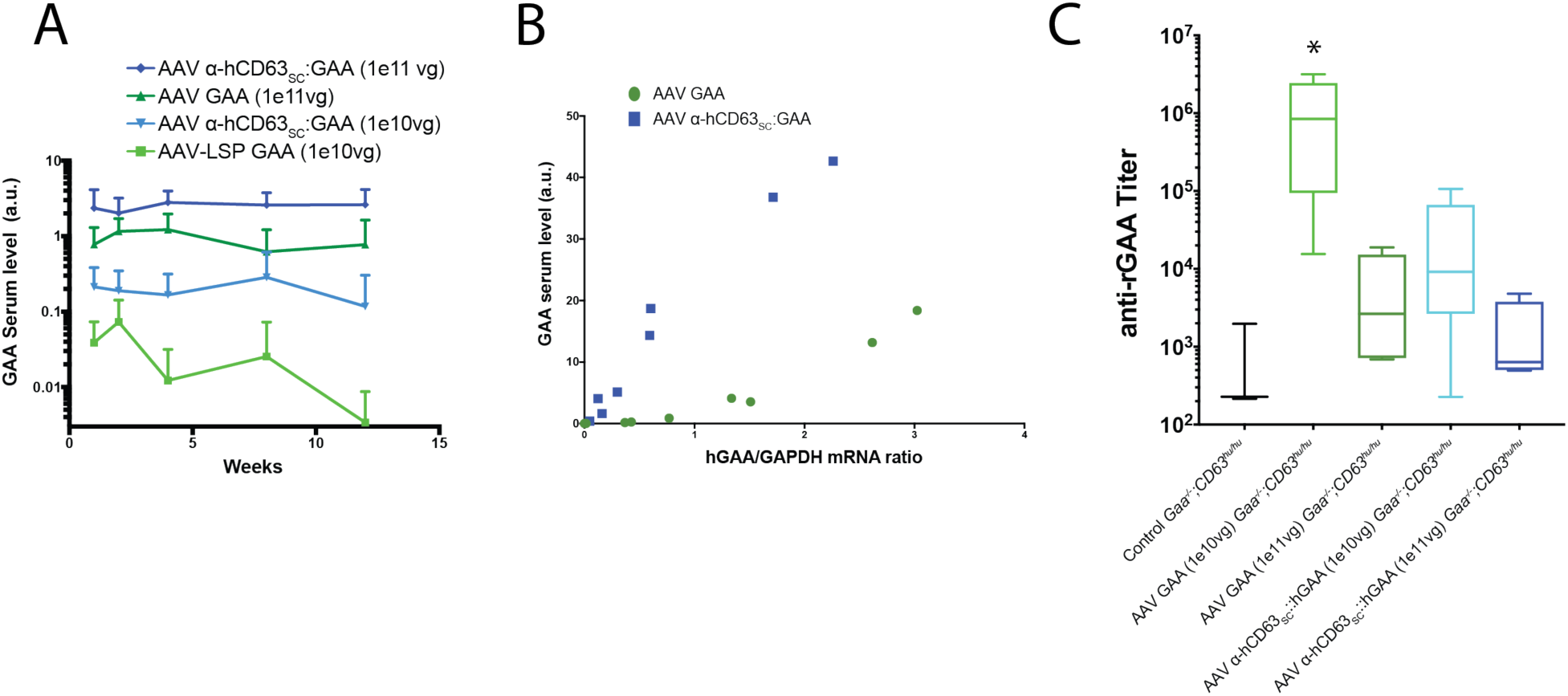
AAV-transduced liver secretes more α-hCD63_SC_:GAA than GAA *in vivo*. (A) α-hCD63_SC_:GAA circulates at higher levels than GAA (n=4 for 1e11vg groups, n=5 for 1e10vg groups). (B) Higher levels of α-hCD63_SC_:GAA are seen in the serum than GAA for a unit of α*-hCD63*_*SC*_:*GAA* or *GAA* mRNA in the liver (each point represents an individual mouse). (C) Higher expression of GAA or α-hCD63_SC_:GAA results in lower anti-GAA titers. Serum IgG titers against rGAA 3 months after infection. Error bars mean are +/− SD. * = p <0.01 vs control group.

Serum titers against GAA performed 3 months after infection revealed that only the 1e10vg AAV GAA group, with a mean titer of 1.18e6, was significantly different versus control (p=0.019) (Figure 2C). The 1e11vg AAV GAA, 1e10vg AAV α-hCD63_SC_:GAA, and 1e11vg AAV α-hCD63_SC_:GAA groups had titers in the 10^4^-10^5^ range, but these did not reach statistical significance versus control mice. These results agree with previous studies showing there is a serum level threshold that must be reached in order to induce tolerance to GAA ^25^.

### AAV-mediated liver depot gene therapy of α-hCD63_SC_:GAA clears muscle glycogen to wild-type levels

To determine if the treatments were effective in reducing lysosomal glycogen in muscle, AAV-infected *Gaa*^*−/−*^;*Cd63*^*hu/hu*^ mice were sacrificed after 3 months, and tissues were assayed for glycogen. Qualitatively, clearance of glycogen to wild-type levels was observed in PAS stained sections of quadriceps after treatment of 1e11vg AAV α-hCD63_SC_:GAA, but not in the 1e11vg AAV GAA treated mice (Figure 3A). Glycogen levels were quantified in muscle tissue lysates (Figure 3B), and in the 1e11vg AAV α-hCD63_SC_:GAA group was reduced to near wild-type levels in heart and all skeletal muscles tested, whereas 1e11vg AAV GAA was not as efficacious, clearing approximately 50% of the glycogen in muscles other than heart. Both soleus (type I myofiber) and extensor digitorum longus (type II myofiber) muscles were cleared of glycogen, with measurements comparable to wild-type levels, in the 1e11vg AAV α-hCD63_SC_:GAA treated group.

**Figure 3.**
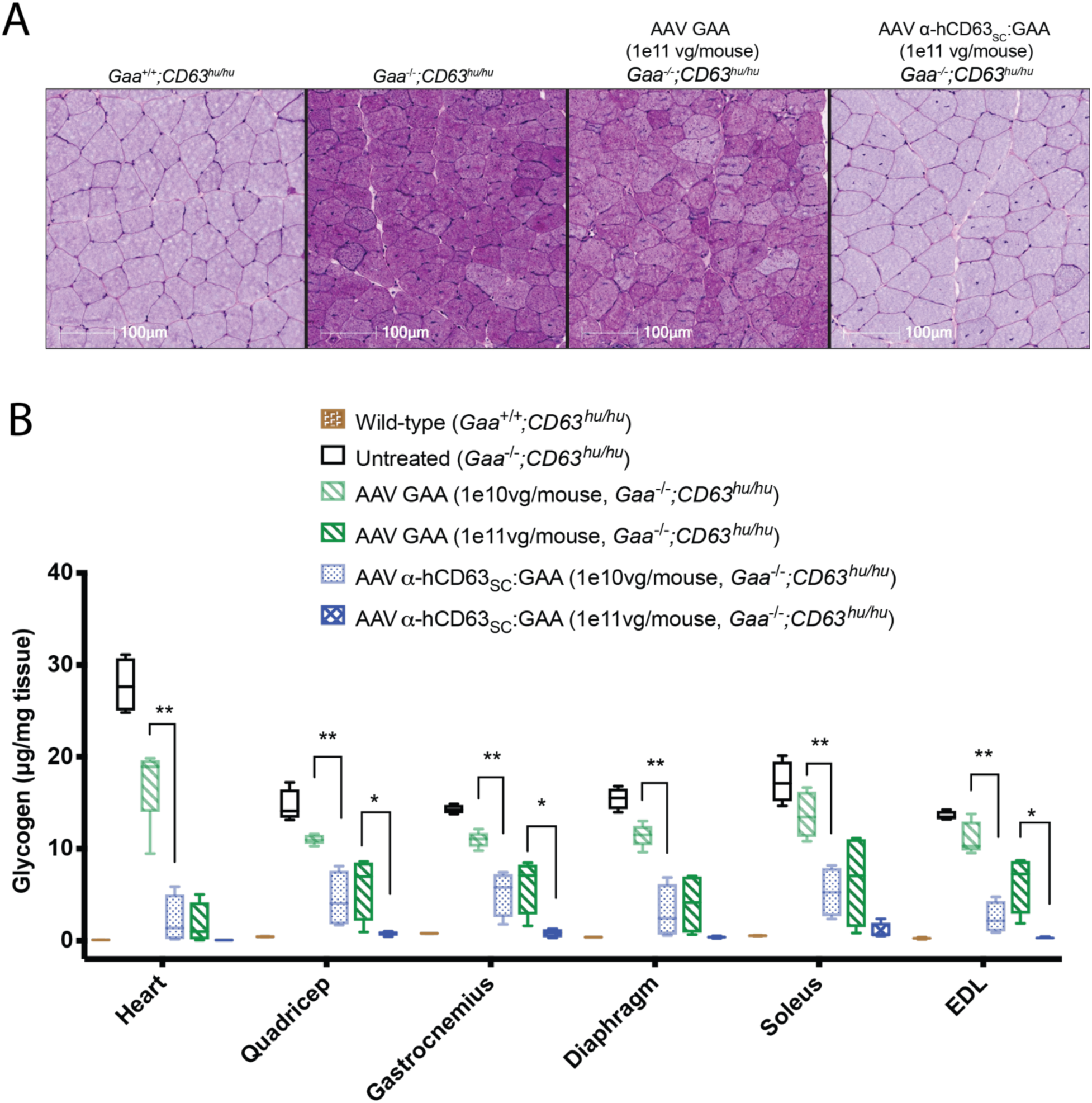
Treatment of PD mice with AAV α-hCD63_SC_:GAA restores glycogen to wild-type levels in key muscles. (A) PAS-H stain for glycogen of 1e11vg treated quadricep muscle sections 3 months after AAV infection show uniform removal of glycogen in the α-hCD63_SC_:GAA group. (B) α-hCD63_SC_:GAA treated mice have lower glycogen levels in skeletal muscle than GAA treated mice at equivalent doses. Normalization of muscle glycogen to wild-type levels is seen at the 1e11vg/mouse dose of α-hCD63_SC_:GAA. Glycogen in muscle tissue lysates were assayed 3 months post AAV-administration. Error bars mean are +/− SD. ** p<0.01 between 1e10vg/mouse groups. * p<0.05 between 1e11vg/mouse groups. n=2 for wild-type, n=4 for treatment groups.

10 months after treatment with the high 1e11vg/mouse dose, CNS tissues were assayed for glycogen removal. Trends of reduction were seen in both AAV α-hCD63_SC_:GAA and AAV GAA treated groups, but the only significant decrease in glycogen levels was in the spinal cord in the AAV α-hCD63_SC_:GAA treated mice (Figure S9).

### *Muscles from mice treated with AAV α*-hCD63_SC_*:GAA showed fewer lysosomes and reduced autophagy*

Lysosomal and autophagosomal accumulation is a hallmark of PD, and effective treatments require their normalization. To visualize lysosomes, sections of quadriceps from the AAV-treated mice and controls were also stained with an anti-LAMP1 antibody and imaged in both confocal (Figure 4A) and wide-field (Figure 4B) microscopy. Quadriceps from *Gaa*^*−/−*^;*Cd63*^*hu/hu*^ mice showed a large expansion of lysosomes (Figure 4C), with lysosomes present throughout the muscle fibers rather than restricted mainly to periphery of the fiber as in wild-type controls (*Gaa^+/+^;Cd63^hu/hu^*) (Figure 4A). Treatment with AAV α-hCD63_SC_:GAA improved lysosomal localization (Figure 4A) and significantly reduced the total lysosomal area (Figure 4B and 4C). In contrast, treatment with AAV GAA showed a trend towards reduced lysosomal area, but the difference from the untreated group was not significant.

**Figure 4.**
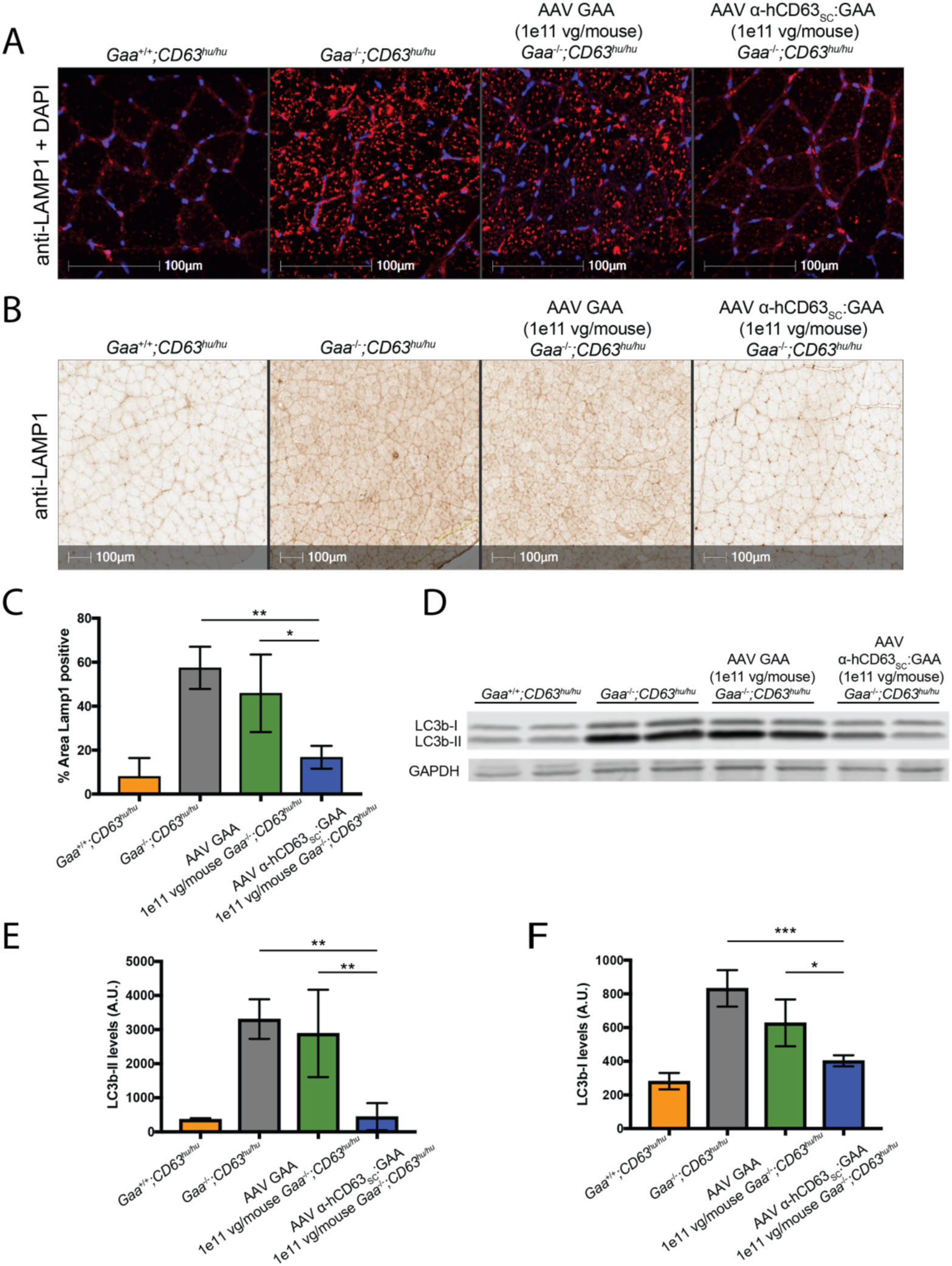
Treatment of PD mice with AAV α-hCD63_SC_:GAA reduces lysosomal area and autophagy. (A) Confocal images and (B) wide-field images of anti-LAMP1 staining to detect lysosomes in section of quadriceps in mice 3 months after administration of AAV GAA or AAV α-hCD63_SC_:GAA show lysosome staining is decreased with scFv:GAA treatment. (C) Quantification of the LAMP-1 positive area in wide-field images. n=2 for wild-type, n=4 for treatment groups. Error bars mean are +/− SD. (D) Autophagy is decreased with scFv:GAA treatment. Representative western blotting for LC3b to monitor autophagy in quadriceps lysate from two mice per group. Each duplicate lane (E) Quantification of LC3b-II and (F) LC3b-I levels in quadriceps lysates. n=2 for wild-type, n=4 for treatment groups. Error bars are mean +/− SD. * = p <0.05, ** = p<0.01, *** = p<0.001.

Increased autophagy is well-documented in PD and in PD mice ^7,26^ and may be a key part of the mechanism of pathogenesis. LC3b is a protein component of the autophagosome and a commonly used marker of autophagy. LC3b-II, a fragment of LC3b that is generated during autophagy, has been previously shown to be elevated in PD mice. To examine autophagy in skeletal muscle, western blotting for LC3b was carried out on quadriceps lysates from the AAV-treated mice and controls (Figure 4D). Compared to wild-type mice (*Gaa+/+;Cd63hu/hu*), quadriceps from *Gaa*^*−/−*^;*Cd63*^*hu/hu*^ mice showed significantly increased levels of both LC3b-I and LC3b-II, with striking increases in LC3b-II (Figure 4D, E, and F), consistent with previous reports ^7,26^. Treatment with AAV α-hCD63_SC_:GAA significantly reduced both LC3b-I and LC3b-II, while treatment with AAV GAA had a much smaller effect (Figure 4D, E, and F).

### *Rotarod and grip strength measurements improved in GAA KO mice following treatment with AAV α*-hCD63_SC_*:GAA*

To assess whether the improved glycogen reduction seen with AAV α-hCD63_SC_:GAA translated into improved muscle function, 2-3 month old *Gaa*^*−/−*^;*Cd63*^*hu/hu*^ mice were treated with either single dose of 1e11 vg/mouse of AAV α-hCD63_SC_:GAA or AAV GAA, or left untreated as controls. Age-matched *Gaa+/+;Cd63hu/hu* served as wild-type controls. The mice were tested on the Rotarod and grip strength apparati one week prior to AAV administration and then monthly for the following 6 months.

Mice treated with AAV α-hCD63_SC_:GAA began showing significantly improved performance compared to untreated mice on both the grip strength and Rotarod tests at 2 and 3 months following AAV administration, respectively (Figure 5 A,B). On both measures, performance of AAV α-hCD63_SC_:GAA treated mice tracked closely with that of WT mice from 3 months of treatment on. Starting at 3 months post-treatment, the performance of mice treated with AAV α-hCD63_SC_:GAA on Rotarod was also significantly better than mice treated with AAV GAA. Mice treated with AAV GAA showed a small trend towards improvement in both measures, but it was not statistically significantly different from untreated mice.

**Figure 5.**
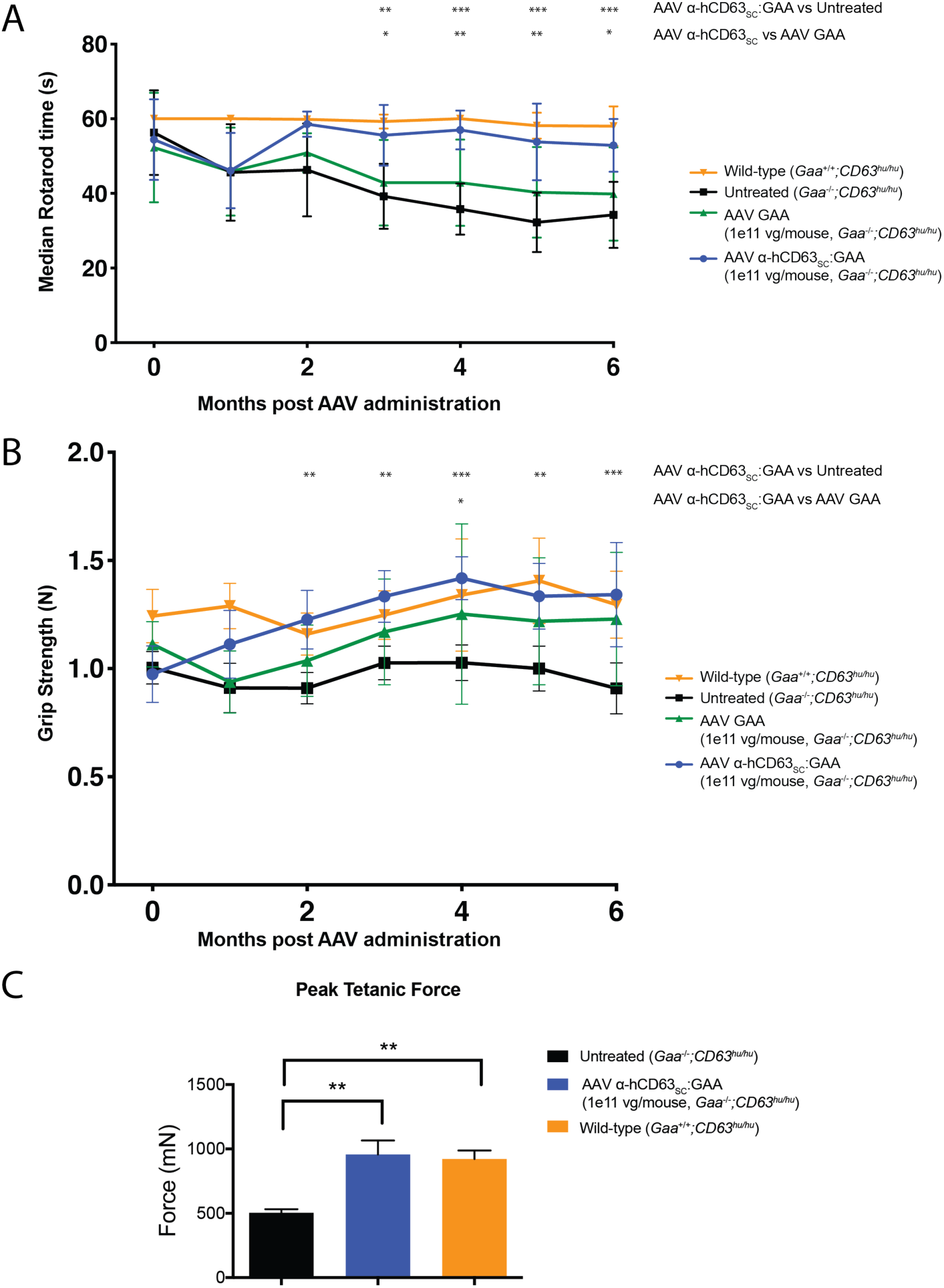
Treatment of PD mice with AAV α-hCD63_SC_:GAA restores muscle function in PD mice. (A) Rotarod test performance of mice treated with either AAV GAA or AAV α-hCD63_SC_:GAA shows recovery of scFv:GAA treated mice within 2 months of AAV administration. (B) Forelimb grip strength measurements show continued strength improvement within 1 month of treatment in α-hCD63_SC_:GAA treated mice. PD mice were treated with 1e11vg AAV α-hCD63_SC_:GAA or 1e11vg AAV GAA or were left untreated, and these groups were compared to wild type mice. (C) Ex vivo peak tetanic force of the tibialis anterior muscle 6 months post AAV-treatment shows a recovery in muscle strength in the α-hCD63_SC_:GAA treated group to comparable to wild-type levels. (A) and (B) Error bars are median +/− SD, n=7 for wild-type, n=11 for treatment groups. (C) Error bars are mean +/− SD, n=4 for untreated and wild-type, n=5 for treatment group. * = p <0.05, ** = p<0.01, *** = p<0.001.

### α-hCD63_huSC_*:GAA has higher glycogen removal at equivalent serum levels compared to GAA*

To determine whether the superior efficacy of α-hCD63_SC_:GAA is due to both improved targeting as well as improved secretion of α-hCD63_SC_:GAA compared to GAA (Figures 2A, 2B, and S8), or to primarily only one of these factors, we used a GAA construct that has several modifications to improve expression and secretion. In this construct, AAV-LSP2 GAA ^27^, the most notable change is a replacement of the endogenous signal peptide with a chymotrysinogen B2 signal peptide. Additionally, in our α-hCD63_SC_:GAA construct we changed the scFv to fully human α-hCD63 variable domains (α-hCD63_huSC_:GAA), as a fully human protein is desirable for possible translation to human trials. Doses of each AAV ranging from 5e11vg/kg to 4e12vg/kg were given to mice to be able to quantify the relationship between residual glycogen and serum level of either α-hCD63_huSC_:GAA or GAA after 2 months of treatment (Figure S10). In mice that received AAV α-hCD63_huSC_:GAA, similar decreases in residual glycogen were seen at approximately a 1 log lower of concentration of α-hCD63_huSC_:GAA in the serum, compared to mice that received AAV-LSP2 GAA and expressed GAA. These results demonstrate that α-hCD63_huSC_:GAA has a higher efficacy due to improved targeting and internalization, and not primarily due to improved secretion.

To better quantify the level of α-hCD63_huSC_:GAA in the serum, we developed an ELISA to detect the scFv portion of the molecule. One month after AAV administration, in mice where the serum level of α-hCD63_huSC_:GAA was >5µg/mL, glycogen levels were normalized to wild-type levels (Figure S11). In comparison, >50µg/mL serum GAA is required for similar levels of glycogen clearance in PD mice ^15^. It is important to note, though, that α-hCD63_huSC_:GAA and GAA may have different rates of clearance from the serum given their different uptake mechanisms, so therefore an equivalent steady-state serum level of α-hCD63_huSC_:GAA and GAA does not necessarily indicate that equivalent numbers of molecules of α-hCD63_huSC_:GAA and GAA are being produced.

## Discussion

We report here targeted ERT, a method for engineering lysosomal enzymes in a manner that enhances their delivery to the tissues where they are needed. Targeted ERT achieves this goal by fusing the enzyme to antibodies that bind transmembrane or cell surface proteins – “effector proteins” – that traffic to the lysosome. The antibody moiety brings the enzyme to the effector protein, which in turn traffics the antibody:enzyme fusion protein to the lysosome. Effector proteins are chosen based on their expression profile which should include or preferably coincide with the tissues affected in the corresponding lysosomal disease. Using PD as an example, we focused on delivery of GAA in PD mice, and chose antibodies against effector proteins that are enriched in cardiac and skeletal muscle, i.e. the target tissues for PD therapy. We demonstrate that targeted delivery of GAA is highly effective and circumvents the limitations of the CI-MPR-mediated delivery (*i.e*. the mechanism on which current ERT relies).

Conceptually, any transmembrane or cell surface protein that is expressed in ‘disease-relevant’ tissues and that internalizes to the lysosome can potentially be used as an effector protein, provided that it is sufficiently abundant and an antibody can be raised against it. In this study, two different types of effector proteins were chosen to test targeted ERT in PD. The first was CD63, a tetraspanin with broad expression across many cell types yet highly expressed in skeletal muscle, and the second was ITGA7, an integrin that is enriched in skeletal and cardiac muscle tissue. Both α-hCD63:GAA and α-ITGA7:GAA enhanced glycogen removal by GAA in PD mice, providing proof-of-concept for targeted ERT. Furthermore, using an scFv version of α-hCD63:GAA with gene therapy as the mode of delivery, we have achieved improved muscle function over the long term in a longitudinal study.

While initial results are promising when the treatment is delivered to young mice, further tests should be performed to determine if a targeted enzyme delivery system could address the decreased efficacy of conventional ERT in older PD mice. Glycogen clearance in mice treated with recombinant GAA late in the course of disease was not sufficient to recover motor function or reverse muscle damage ^8^. The role of insufficient CI-MPR mediated delivery, while significant, may play a much smaller role than the inability of muscle to repair severely damaged fibers in aged mice ^28^.

Additionally, recent evidence suggests a role for glycogen accumulation in tissues other than cardiac and skeletal muscle in the pathology of PD. Abnormalities in the neuromuscular junction have been linked with glycogen deposits in spinal motor neurons in mice ^4^, and infantile-onset patients show secondary symptoms indicative of neural involvement ^3,29^. Studies normalizing glycogen levels solely in the brain and CNS in PD mice (using AAV gene therapy) have shown correction of some neuromuscular phenotypes even in the absence of improvement in cardiac or skeletal muscle fibers, demonstrating that a full reversion of clinical phenotypes in PD ERT will require multisystem delivery ^30,31^. While we and others have seen a trend or partial reduction in CNS glycogen storage (Supplementary Figure S9) ^27^, it remains unknown whether this reduction will be clinically significant in PD patients. To maximize enzyme delivery to affected tissue types, an effector like CD63, which – in addition to cardiac and skeletal muscles – is also expressed in the CNS, would likely be more appropriate than more tissue-restricted internalizers like ITGA7. Therefore, adding blood-brain barrier crossing functionality to neuronal lysosome targeting in a format such as a bispecific antibody may be able to improve overall efficacy ^32,33^. Alternatively, AAV may be delivered via an intrathecal or intracerebroventricular route as well as intravenously to treat both the CNS and skeletal muscle ^29,30^.

ERT for other lysosomal diseases, such as rGALNS for mucopolysaccharidosis VIA ^34^ and rARSB for mucopolysaccharidosis VI ^35^, also have poor delivery to diseased non-CNS organs partially due to the limitations of CI-MPR ^36^. Enzyme delivery to address the skeletal phenotypes in mucopolysaccharidoses I-VII remains a challenge due to the virtually nonexistent delivery to these tissues ^37^. Antibody targeting of these enzymes in a similar manner to that presented here using internalizers enriched in growth plate or articular cartilage cells may be able to enhance enzyme delivery and correction of the skeletal phenotypes.

The approach to clinical development of targeted ERT will be determined by the therapeutic format, as targeted ERT can be administered as either a purified protein or a gene therapy. Infusion of a purified protein should be considered due to its maturity as a therapeutic modality, whereas a gene therapy approach may offer several potential advantages over a protein-based ERT. One potential advantage of the AAV approach is that it alleviates the need for every other week infusions of ERT. However, liver-targeted AAV-based ERT is unlikely to lead to life-long expression of the transgene due to hepatocyte turnover, especially in younger patients ^38^.

Furthermore, neutralizing antibodies (nAbs) against the AAV capsids may not only exclude patients from treatment eligibility but may further prevent re-dosing AAV therapies, although immunomodulation and tolerization approaches are currently under study that may show a path forward to overcome nAbs against AAV capsids ^39^. Another potential advantage of the AAV approach is that it may help overcome nAbs against the replacement enzyme. An immune response to rhGAA is known to occur in a significant number of PD patients and often requires significant clinical intervention ^40^. Immunogenicity of the replacement enzyme still remains a major concern with targeted ERT, and immunogenicity may even be enhanced if the antibody directs the fusion protein to antigen presenting cells or due to creation of possible neoepitopes in the scFv or junctions of the fusion protein. Liver depot gene therapy has been shown to produce immune tolerization in mice, and promising evidence for the ability of liver depot gene therapy to reverse pre-existing anti-drug antibodies has been shown in mice ^38,39^. Clinical trials to test whether this is true in humans have already begun. Finally, it should be noted that the modularity and interchangeability of the targeting antibody is an inherent advantage to this platform technology over delivery technologies that utilize a single binding domain/receptor interaction. Swapping (or even combining) targeting antibodies to different protein targets to tune delivery to desired cell types, while avoiding other cell types, will be a significant advance in the field of enzyme replacement therapy.

## Methods

### Plasmid backbones for protein production, HDD, and AAV

Antibody-GAA plasmid constructs were created by isothermal assembly^41^. Antibody:GAA fusion constructs were cloned with a human IgG4 antibody backbone, an amino acid linker comprising GGGGS, and amino acids 70-952 of human GAA. Full-length human GAA cDNA (amino acids 1-952) plasmids were used as a comparator. Protein production plasmids used a CMV promoter, HDD plasmids used an ubiquitin C promoter, and AAV plasmids used a liver-specific transthyretin promoter with a liver enhancer from *Serpina1* ^42^. scFv fusions were cloned with domains in the following order: Vh, 3x glycine-serine linker, Vk, 1x glycine-serine linker, amino acids 70-952 of human GAA.

### Antibody variable domains

Anti-mouse CD63 antibodies and their fusions used the NVG-2 rat anti-mouse CD63 variable domains (a kind gift from Drs. Miyasaka and Hayasaka, Osaka University) ^43^. Anti-human CD63 antibodies and their fusions used the H5C6 mouse anti-human CD63 variable domains (Iowa Developmental Studies Hybridoma Bank). Anti-mouse integrin alpha-7 antibodies and their fusions used the variable domains of mouse anti-mouse integrin alpha-7 non-blocking antibody 3C12 ^44^.

### Protein purification of antibody-GAA constructs

Antibody:GAA constructs were transfected into CHO cells and purified by protein A or protein L chromatography. Briefly, CHO supernatants were flowed through columns packed with Protein L or Protein A (GE Healthcare) in 50mM Tris, pH7.5, 150mM NaCl. Columns were washed with 50mM Tris, pH7.5, 500mM NaCl. Proteins were eluted with IgG Elution buffer at pH 2.8 (Pierce) and neutralized with 1M Tris, pH 8.5. The protein solution was buffer exchanged to 20mM Sodium Phosphate, 150mM NaCl, pH 6.2 by dialysis. Size-exclusion chromatography on Superose 6 PGS was used to remove high molecular weight contaminants. Final product was stored in 20mM Sodium Phosphate, 150mM NaCl, pH 6.2.

### In vitro internalization of antibody-GAA constructs

Primary adult human myoblasts (Lonza CC-2561), infantile Pompe fibroblast lines GM20089 and GM20090 (Coriell), C2C12 myoblasts, and CRISPR-modified HEK293 cells were used to test internalization of antibody-GAA constructs. GM20089 fibroblasts are missing exon 18 of GAA, and GM20091 fibroblasts are a compound heterozygote of Q58X and del525T. Both lines are deficient in processing of GAA pre-protein to its mature lysosomal form. Antibody:GAA proteins were incubated with semi-confluent cells overnight, washed extensively with PBS, and then lysed with either RIPA buffer for protein analyses or sodium acetate lysis buffer (0.2mM sodium acetate, 0.4mM potassium chloride, 0.5% NP-40, pH 4.3). Cells were incubated with leupeptin or E-64 (Sigma-Aldrich L9783, E3132) in experiments testing GAA intracellular processing and cleavage. For pulse-chase experiments, cells were incubated overnight with a dose of 50nM α-hCD63:GAA in the media. After incubation, cells were washed once with PBS and given fresh media. Media was changed every three days over the course of the experiment.

### Generation of CD63-deficient HEK293 cells by CRISPR

To generate *CD63-deficient* HEK293 cells, cells were co-transfected using LT1 reagent and protocol (Mirus Bio, MIR2306) with a plasmid containing Cas9+NLS under the CMV promoter and a plasmid containing the guide RNA CACTCACGCAAAAGGCCAGCAGG under the U6 promoter. Pooled transfected cells were then trypsinized, stained with α-hCD63 antibody H5C6 (Iowa Developmental Studies Hydridoma Bank) and goat anti-mouse Alexa647 (Invitrogen) were then subject to FACs sorting. The cell population negative for anti-hCD63 staining was collected and expanded. The pool was then further confirmed to have uniform negative staining for CD63 by flow cytometry using the same antibodies.

### GAA knockout mouse models

The GAA^6neo/6neo^ mouse line ^23^ was obtained from The Jackson Laboratory.

*Cd63* humanized, *Gaa* homozygous-null (*Cd63*hu/hu *Gaa*^−/-^) mice were generated using the VelociGene™ method ^45,46^. Nucleotide coordinates correspond to those in mouse genome Ensembl release 98 (September 2019) are provided to facilitate a precise description of the alleles generated. To begin, an allele with fully humanized Cd63 coding sequence was generated via recombineering technology. Mouse genome sequence with the coordinates 128911372-128912754 on Chromosome 10, which is contained with BAC RP23-122I15, was replaced with the Human genome coordinates 55725292-55727639 from Chromosome 12. This replaces the mouse CD63 exons 2-7 (ENSMUSE00000150030-ENSMUSE00001408024) with the orthologous human CD63 exons, part of the 1^st^ intron, intervening introns, and 3’ non-coding sequences. Exon 1 (ENSMUSE00000150024) was not replaced as it encodes protein sequence 100% identical to human. A self-deleting, fLoxed hygromycin resistance cassette was placed in the intron following human coding exon 5 (ENSE00003590609) between coordinates 12:55729738-55728596. The self-deleting cassette contained the hygromycin resistance coding sequence under the control of the human *UBC* promoter and an Em7 promoter for growth in *E. coli*, followed by a poly(A) signal from mouse *Pgk1.* For deletion in the F0 male germline, the cassette also contained Cre coding sequence controlled by the mouse *Prm1* promoter and an SV40 poly(A) signal. The Cre coding sequence was interrupted by a synthetic intron, to prevent expression in bacteria. The resulting targeting construct had 125 and 84 Kb homology arms, respectively and contained a hybrid *Cd63/CD63* allele encoding protein 100% identical to human CD63. A GenBank file with the sequence of the targeting construct is supplied as supplementary materials. The targeting construct was then linearized with *NotI*, and electroporated into 50% C57BL/6NTac/50% 129S6/SvEvTac embryonic stem cells to generate allele VG7232 (*Cd63*^+/hu-HYG^).

To generate allele VG4323 (*Gaa*^+/NEO^), a cassette consisting of a lacZ reporter and a self-deleting, fLoxed neomycin resistance cassette (as above, but with neomycin resistance in place of hygromycin resistance) was inserted into mouse BAC bMQ-208N21, replacing the mouse genome coordinates 119270136-119285130 on Chromosome 11 which include part of exon 2 (ENSMUSE00000151834), exons 3-19, and part of exon 20 (ENSMUSE00000364002). This yielded a final targeting construct with 11 and 18 Kb homology arms, respectively. The cassette was inserted with linker such that lacZ coding sequence was directly in frame with the initiating Methionine in *Gaa* while the complete open reading frame of *Gaa* was replaced. A GenBank file with the sequence of the targeting construct is supplied as supplementary materials. This targeting vector was then linearized with *NotI* and electroporated into *Cd63*^+/hu-HYG^ embryonic stem cells. Following microinjection, F_0_ offspring were bred to wildtype C57BL/6NTac and genotyped for alleles VG7233 (*Cd63*^+/hu^). and VG6533 (*Gaa*^+/-^), resulting from male germline deletion of the hygromycin and neomycin cassettes, respectively. Animals were then intercrossed to generate *Cd63*^hu/hu^;*Gaa*^−/-^ cohorts used for subsequent experiments.

### AAV production and in vivo transduction

Recombinant adeno-associated virus 8 (AAV2/8) was produced in HEK 293 cells. Cells were transfected with three plasmids encoding adenovirus helper genes, AAV8 rep and cap genes, and recombinant AAV genomes containing transgenes flanked by AAV2 inverted terminal repeats (ITRs). On day 5, cells and medium were collected on day 3, centrifuged, and processed for AAV purification. Cell pellets were lysed by freeze-thaw and cleared by centrifugation. Processed cell lysates and medium were overlayed onto iodixanol gradients columns and centrifuged in an ultracentrifuge. Virus fractions were removed from the interface between the 40 and 60 percent iodixanol solutions and exchanged into 1xPBS using desalting columns. AAV viral genomes (vg) were quantified by qPCR using TaqMan oligos targeting the ITR. A standard curve was generated using serial dilutions of virus with a known concentration. AAVs were diluted in PBS + 0.001% F-68 Pluronic immediately prior to injection.

### Muscle performance assays

AAV-treated mice were subjected to monthly Rotarod and forelimb grip strength measurements. The Rotarod (IITC Life Sciences) was programmed to accelerate at 0.5rpm/sec for 60 sec, and time to fall was recorded. For the grip strength testing, mice were placed on a flat wire mesh, and the mesh was inverted. Time to fall was recorded with a maximum of 60 sec. Forelimb grip strength was measured using a force meter (Columbus Instruments, Ohio, USA). All tests were performed in triplicate at each time point.

### Ex vivo muscle physiology

To determine whether AAV α-hCD63_SC_:GAA treatment affected TA muscle function, ex vivo force was measured 6 months post dosing as previously described ^47^. Briefly, TA muscles were removed from anesthetized mice from Pompe untreated, AAV α-hCD63_SC_:GAA, and GAA wild-type mice. Maximal twitch force and peak isometric tetanic force were measured ex vivo immediately after dissection.

### Tissue collection and glycogen measurements

Tissues were dissected from mice immediately after CO_2_ asphyxiation, snap frozen in liquid nitrogen, and stored at −80ºC. Tissues were lysed on a benchtop homogenizer with stainless steel beads in distilled water for glycogen measurements, or RIPA buffer for protein analyses. Glycogen analysis lysates were boiled and centrifuged to clear debris. Glycogen measurements were performed fluorometrically using a commercial kit according to manufacturer’s instructions (Biovision K646).

### Serum titers

Serum from tail or cheek bled mice was collected using serum separator tubes (BD Biosciences, 365967) High protein-binding 96-well plates were coated with 2µg/mL GAA overnight. Plates were blocked with 0.5% BSA. Serial dilutions of mouse serum were incubated overnight on the plates to bind to GAA. Total mouse IgG against GAA was measured using HRP-goat anti-mouse IgG (subclasses 1 + 2a + 2b + 3) (Jackson Immunoresearch, 115-035-164) and a colorimetric kit (BD Biosciences, 555214). Reactions were stopped using 1N phosphoric acid. Absorbances were read at 450nm. The dilution curves were fit to sigmoidal curves, and antibody titers were calculated.

### Histology

Mouse tissues were embedded immediately after dissection in optimal cutting temperature (Tissue-Tek O.C.T.) chilled in a bath of isopentane chilled in a bath of liquid nitrogen. Blocks were stored at −80°C until they were cut at 12µm and mounted onto slides. Sectioning and PAS-H staining was performed by Histoserv (Germantown, MD). For antibody staining, slides were post-fixed in 4% paraformaldehyde (Electron Microscopy Services 15710), blocked in eBioscience Blocking Buffer (LifeTech Thermo 00-4953-54) followed by staining with anti-Lamp1 1D4B (Abcam ab25245). For fluorescent imaging, slides were then stained with goat anti-Rat Alexa555 (LifeTech Thermo A21434) and mounted in Fluoromount-G with DAPI (LifeTech Thermo 00-4959-52) and imaged using a Zeiss LSM 710. For visible light imaging, slides were quenched with 30% hydrogen peroxide prior to blocking as above. After the primary antibody, then treated with biotinlated donkey anti-rat IgG (LifeTech Thermo, A18743) followed by DAB staining using ABC kit (Vector Labs PK-6100) and DAB (Sigma D5637). Slides were then dehydrated in alcohols, cleared in xylene, coverslipped and imaged on an Aperio slide scanner. Images were quantified using Halo software.

### Western blots

Cell and tissue lysates were prepared by lysis in RIPA buffer (Millipore, 20-188) with protease inhibitors (Thermo 1861282). Tissue lysates were homogenized using a bead homogenizer (MPBio). Cells or tissue lysates were run on SDS page gels using the Novex system (LifeTech Thermo, XPO4200BOX, LC2675, LC3675, LC2676). Gels were transferred to either nitrocellulose membrane (Li-Cor 926-31090) or low-fluorescence PVDF membrane (Li-Cor IPFL07810) followed by blocking with Odyssey blocking buffer (Li-Cor 927-500000) or 5% milk (LabScientific M0841) in TBS-T and staining with antibodies against either GAA (Abcam, ab137068), anti-LC3B (Sigma L7543), or anti-GAPDH (Abcam ab9484) and the appropriate secondary (Li-Cor 926-32213 or 925-68070). Blots were imaged using a Li-Cor Odyssey CLx. For Figure 1E only, blots were imaged using the chemiluminescent method. Briefly, membranes were incubated with antibodies against GAA (Abcam ab113021) and detected with anti-rabbit or anti-human HRP-conjugated secondary antibodies (Promega W4011 and W4031). Blots were developed using ECL (ThermoScientific 32106) and imaged on X-ray film (Carestream 1651454).

### GAA enzymatic assay

Proteins or cell lysates were assayed for GAA activity using the fluorogenic substrate 4-methylumbelliferyl-alpha-D-glucopyranoside. 4-methylumbelliferone was used as a standard. Fluorescence was measured on a plate reader. GAA activity was calculated as nmol of 4-methylumbelliferyl-alpha-D-glucopyranoside hydrolyzed per hour per nmol of protein. Recombinant human GAA (R&D Biosystems) was used as a standard.

### Statistical Analyses

All statistical tests and analyses were performed in Graphpad Prism, versions 7 or 8. Statistical significance was set at p < 0.05. To compare groups, one-way ANOVAs were performed with post-hoc Dunnett’s multiple comparison tests performed versus controls to determine significance. For grip strength and rotarod test data, statistical significance was determined using 2-way ANOVA with Dunnett’s and Tukey’s multiple comparison tests.

### Data Availability

Regeneron materials described in this manuscript may be available to qualified academic researchers upon request through our portal (https://regeneron.envisionpharma.com/vt_regeneron/). In certain circumstances in which we are unable to provide a particular proprietary reagent, an alternative molecule may be provided that behaves in a similar manner. Additional information about how we share our materials can be obtained by contacting Regeneron’s preclinical collaborations email address.

## Authorship Contributions

A.D.B., C.J.S., A.J.M., A.N.E. and K.D.C. designed the studies. A.D.B., P.T.C., M.P., M.S.B., C.P., and N.L. performed *in vitro* studies. A.D.B., P.T.C., N.A.A., L.M., K.D.C., and L.W. performed the mouse *in vivo* studies. A.M., S.M.T., and C.A.K. designed and produced AAVs. S.B. and A.O.M. designed and produced the transgenic mouse models. P.B. and N.W.G. phenotyped the mouse models. A.D.B., A.J.M., A.N.E., and K.D.C. wrote and edited the manuscript.

