## Supplementary Figures and Methods for "Targeted delivery of acid alpha-glucosidase corrects skeletal muscle phenotypes in Pompe disease mice"

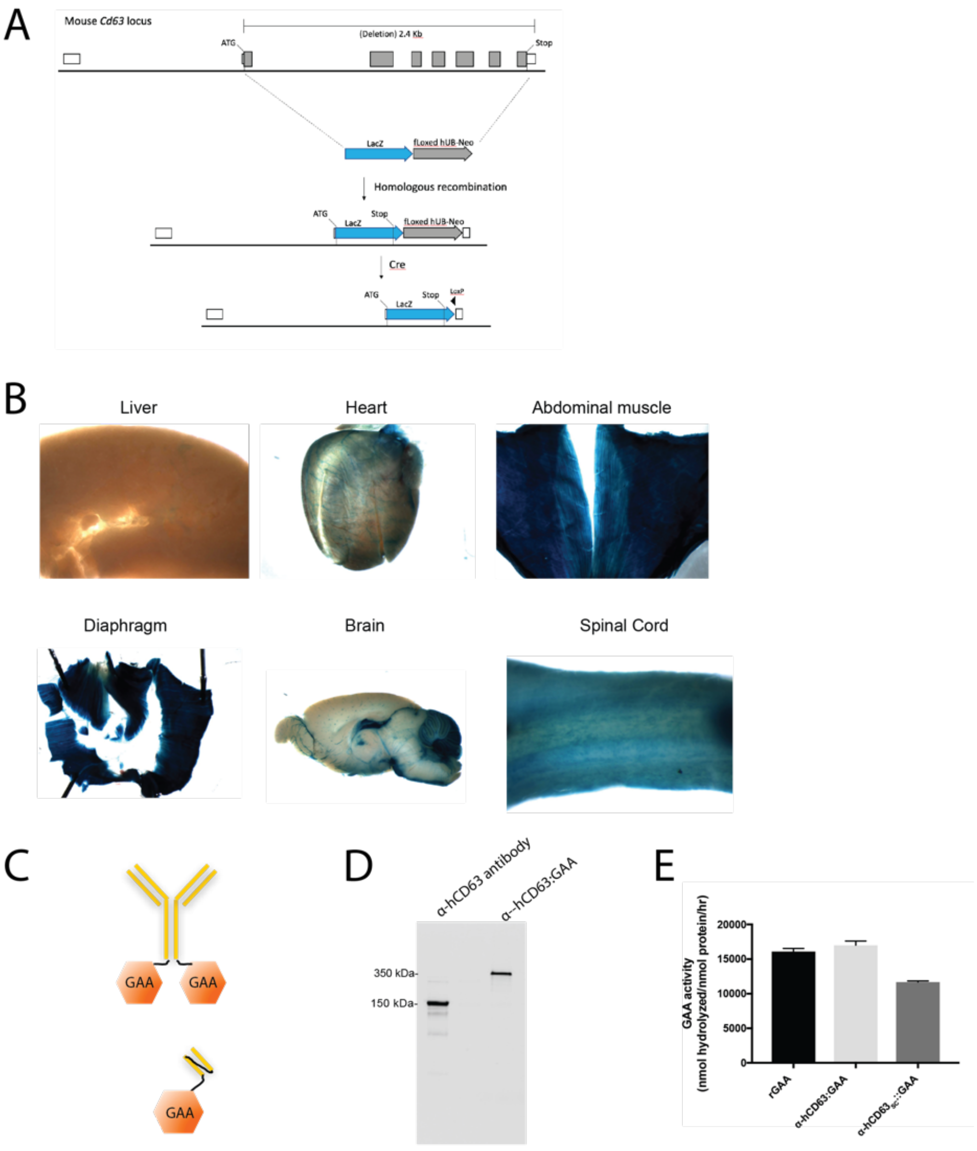

**Supplementary Figure 1.** CD63 is a suitable effector for targeted ERT in mouse PD. (A) Design of a *Cd63*$^{lacZ/lacZ}$ reporter mouse allele. The entire coding sequence of *Cd63* is deleted and replaced with *lacZ.* (B) LacZ staining of sections from liver, heart, abdominal muscle, and diaphragm tissues taken from a *Cd63*$^{lacZ/lacZ}$ reporter mouse verifies expression of CD63 in different skeletal muscles. (C) Schematic of antibody:GAA, a full-length IgG4 or an scFv fused to GAA (amino acids 70-952) with a glycine-serine linker in between the two domains. (D) Western blot probed for hIgG of supernatants of CHO cells transfected to produce α-hCD63 antibody alone or α-hCD63:GAA fusion. No cleavage products of the secreted antibody:GAA protein are visible in the supernatant. (E) GAA enzymatic activities of purified protein lots of GAA, α-hCD63:GAA, and α-hCD63$_{SC}$:GAA.

| Glyco-peptides [a] | Mass Calculated | | | Glycan Abbreviation | Glycan Structure | Relative Abundance (%) |
|---|---|---|---|---|---|---|
| | GlycoPep Mass | Peptide Mass | Sugar Chain Mass | | | |
| Asn849 (hGAA 470) (GVFIT N E TGQPLIGK ) | 3777.62 | 1572.85 | 2204.77 | G2S2 | 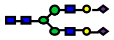 | **18.3** |
| | 3486.52 | | 1913.67 | G2S | 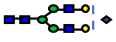 | **34.0** |
| | 3195.43 | | 1622.58 | G2 | 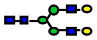 | **16.8** |
| | 3121.39 | | 1548.54 | G1S-GlcNAc | 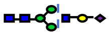 | **3.9** |
| | 2951.33 | | 1378.47 | Man6 | 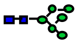 | **9.3** |
| | 2830.30 | | 1257.45 | G1-GlcNAc | 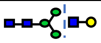 | **5.0** |
| | 2789.27 | | 1216.42 | Man5 | 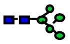 | **10.9** |
| | 2668.25 | | 1095.40 | G0-GlcNAc | 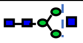 | **1.8** |
| Asn769 (hGAA 390) (QVVE N M TR) | 3180.25 | 975.48 | 2204.77 | G2S2 | 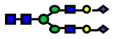 | **14.2** |
| | 2889.15 | | 1913.67 | G2S | 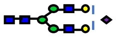 | **27.0** |
| | 2524.02 | | 1548.54 | G1S-GlcNAc | 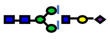 | **4.2** |
| | 2232.93 | | 1257.45 | G1-GlcNAc | 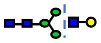 | **3.4** |
| | 2191.90 | | 1216.42 | Man5 | 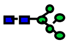 | **51.2** |
| Asn293 (hIgG4) (EEQF N ST YR) | 2941.14 | 1172.51 | 1768.63 | G2F | 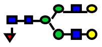 | **8.4** |
| | 2779.10 | | 1606.59 | G1F | 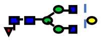 | **40.1** |
| | 2633.04 | | 1460.53 | G1 | 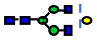 | **1.6** |
| | 2617.04 | | 1444.53 | G0F | 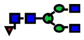 | **47.0** |
| | 2413.97 | | 1241.46 | G0F-GlcNAc | 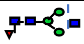 | **2.8** |
| Asn1031 (hGAA 652) showed too low abudant glycopeptides to quar titate | | | | | | |

| Glyco-peptides | Mass Calculated | | | Glycan Abbreviation | Glycan Structure | Relative Abundance (%) |
|---|---|---|---|---|---|---|
| | GlycoPep Mass | Peptide Mass | Sugar Chain Mass | | | |
| **Asn519 (hGAA 140)** (LE N LSSSE MGYTATL TR) | 3566.49 | 1871.89 | 1694.60 | G1FS-GlcNAc | 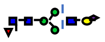 | **8.7** |
| | 3640.53 | | 1768.63 | G2F | 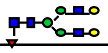 | **12.0** |
| | 3931.63 | | 2059.73 | G2FS | 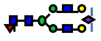 | **48.9** |
| | 4222.72 | | 2350.82 | G2FS2 | 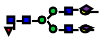 | **30.4** |
| **Asn1304 (hGAA 925)** (VTVLGV ATAPQQV LSNGVPV S N FTYSPD TK) | 5804.57 | 3088.61 | 2715.96 | G3FS2 | 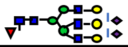 | **12.6** |
| | 5439.44 | | 2350.83 | G2FS2 | 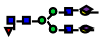 | **40.7** |
| | 5148.34 | | 2059.73 | G2FS | 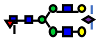 | **44.2** |
| | 5164.35 | | 2075.74 | Hybrid | | **2.4** |
| **Asn612 (hGAA 233)** (VLL N TTV APLFFAL) | 6647.19 | 5106.68 | 1540.52 | Man7 | 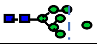 | **33.5** |
| | 6485.14 | | 1378.47 | Man6 | 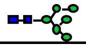 | **60.0** |
| | 6323.09 | | 1216.42 | Man5 | 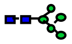 | **6.5** |
| **Asn1261 (hGAA 882)** (N NTIVNE LVR) | 3886.60 | 1170.64 | 2715.96 | G3FS2 | | **4.9** |
| | 3521.46 | | 2350.83 | G2FS2 | 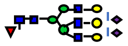 | **36.1** |
| | 3230.37 | | 2059.73 | G2FS | 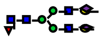 | **43.1** |
| | 2939.27 | | 1768.64 | G2F | 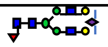 | **12.0** |
| | 2574.14 | | 1403.51 | G1F-GlcNAc | 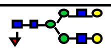 | **3.7** |

Monosaccharide symbols: fucose (fuc); GlcNAc (NAG); mannose (man); galactose (gal); sialic acid

**Supplementary Figure S2**. Tryptic mapping glycan analysis of α-hCD63:GAA protein detects no M6P residues on GAA’s 7 or the α-hCD63 antibody’s 1 N-glycosylation Asn residues.

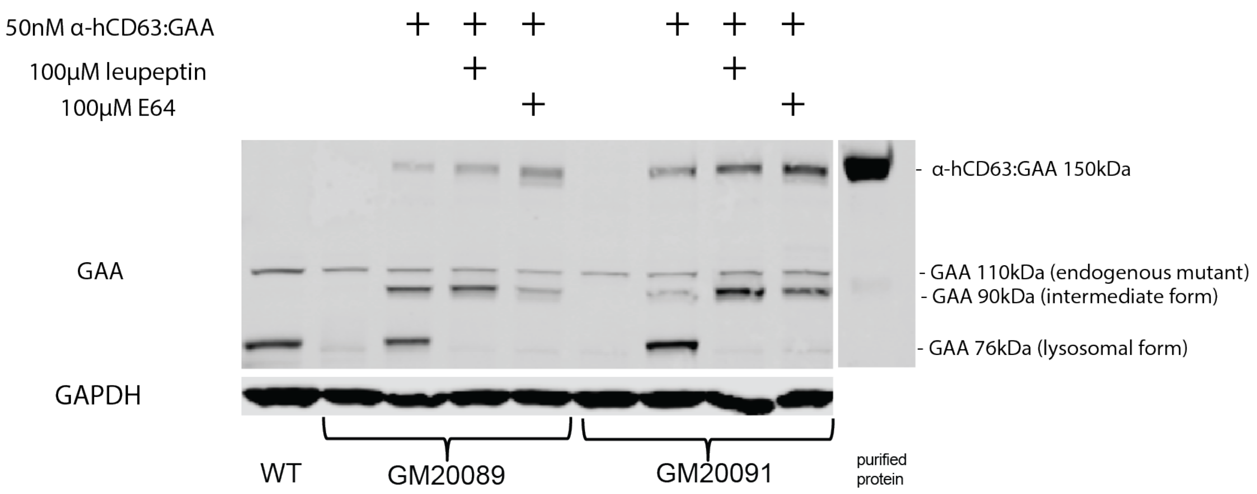

**Supplementary Figure S3.** Antibody:GAA intracellular processing to the lysosomal form of GAA is dependent on cathepsins. Wild-type and two infantile Pompe fibroblast lines (GM20089 and GM20091, which lack the lysosomal GAA forms) were incubated with 50nM α-hCD63:GAA. Either leupeptin or E64 was added to the cells to block cathepsins. Cathepsin inhibition by leupeptin or E64 prevented cathepsin-cleavage of the antibody:GAA construct to the 76kDa lysosomal form of GAA, but did not prevent the processing to the ~90kDa form of GAA.

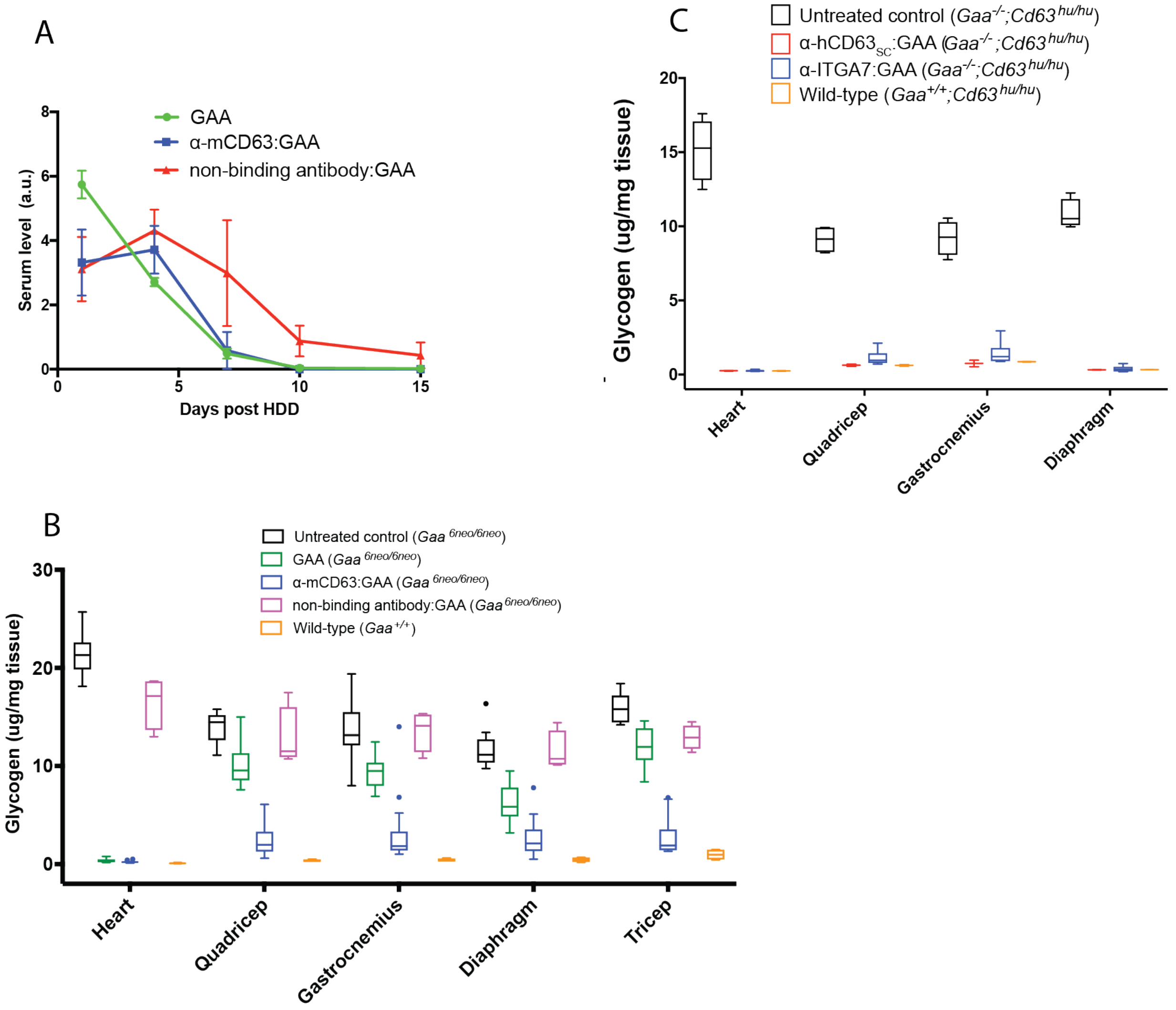

**Supplementary Figure S4**. *In vivo* glycogen reduction is achieved in two mouse models of PD by short-term liver depot gene therapy. (A,B) *Gaa*$^{6neo/6neo}$ mice were dosed with plasmids encoding α-mCD63:GAA, GAA, or a isotype control antibody:GAA by HDD. Serum levels were quantified by western blot, whereas glycogen in tissue lysates was measured 3 weeks post-HDD. (A) Both fusion proteins had similar levels in serum and became undetectable 2 weeks after HDD. (B) α-mCD63:GAA clears glycogen more efficiently than GAA, and is also efficacious in reducing glycogen in tissues where GAA is not effective. The non-binding antibody:GAA negative control does not significantly decrease glycogen levels in muscle. (C) α-hCD63:GAA is effective in treating PD in humanized PD mice. Mice homozygous-null for *Gaa* and humanized for *Cd63* (*Gaa*$^{-/-}$;*Cd63*$^{hu/hu}$) were dosed with plasmids encoding an scFv:GAA format of α-hCD63:GAA or a full-length IgG4:GAA format of α-ITGA7:GAA by HDD. Tissue glycogen levels were measured 3 weeks post-HDD. α-ITGA7:GAA was as effective in removing glycogen as α-hCD63:GAA.

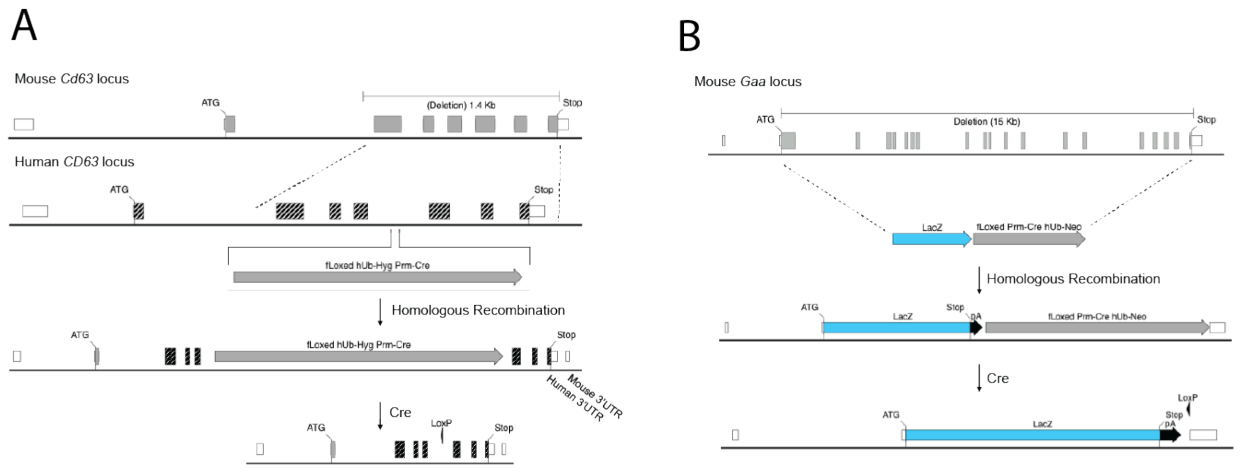

**Supplementary Figure S5.** Design of a PD mouse model with humanized CD63. (A) An allele with humanized *Cd63* coding sequence was generated via recombineering technology. Exons 2-7, as well as part of the intron preceding coding exon 2, and the *Cd63* 3' noncoding region in the mouse locus was replaced with orthologous human *CD63* intronic, coding, and 3' noncoding sequences. Mouse coding exon 1 was not humanized because it encodes protein sequence 100% identical to human. A self-deleting fLoxed hygromycin resistance cassette was placed in the intron following human coding exon 4. The resulting hybrid *Cd63/CD63* allele encodes protein 100% identical to human CD63. (B) A *Gaa* lacZ replacement allele was generated via recombineering technology. A cassette consisting of a lacZ reporter and a self-deleting fLoxed neomycin resistance module replaces the entire *Gaa* coding sequence.

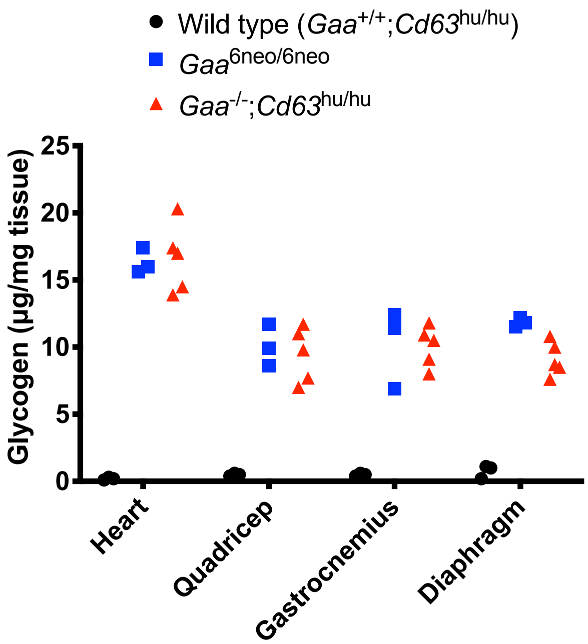

**Supplementary Figure S6**. Glycogen storage is equivalent in *Gaa*$^{6neo/6neo}$ and *Gaa*$^{-/-}$;*Cd63*$^{hu/hu}$ mice. Glycogen content of heart, quadricep, gastrocnemius, and diaphragm tissue lysates from 2 month old mice were measured. n=3 for wild-type and *Gaa*$^{6neo/6neo}$, n=5 for *Gaa*$^{-/-}$;*Cd63*$^{hu/hu}$.

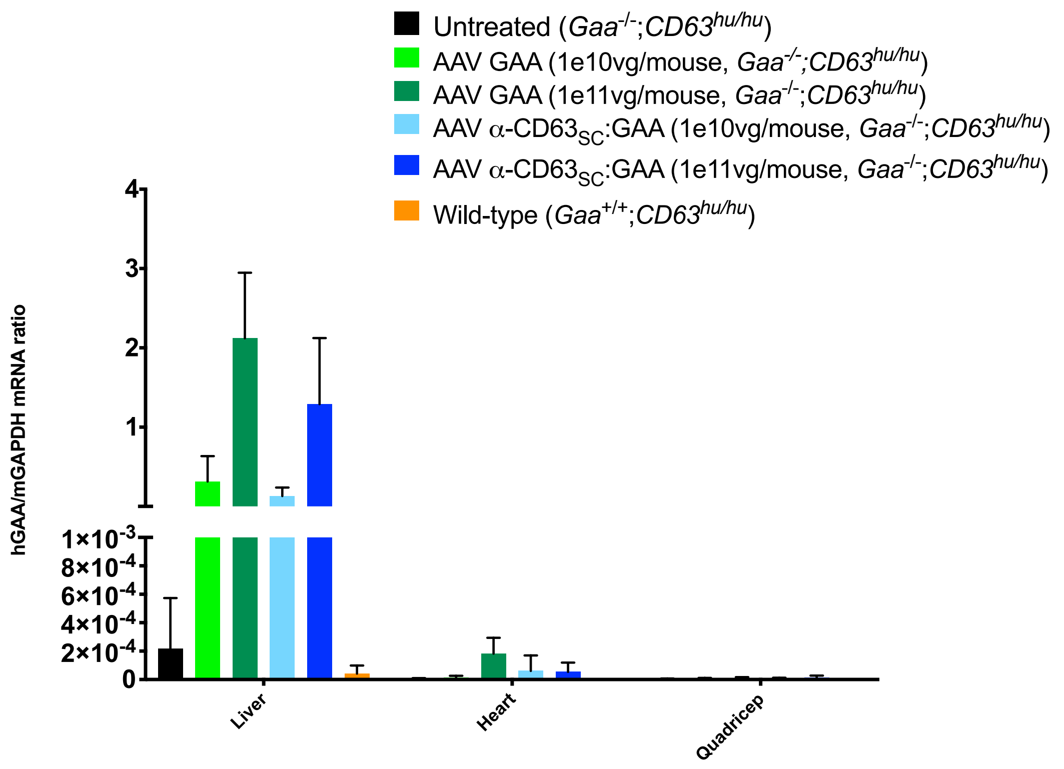

**Supplementary Figure S7.** Real-time PCR quantifications of expression in liver, heart, and quadriceps lysates 3 months after infection shows that expression of α-hCD63$_{SC}$:GAA and GAA is highest in the liver. mRNA levels of α-hCD63$_{SC}$:GAA and GAA do not differ significantly.

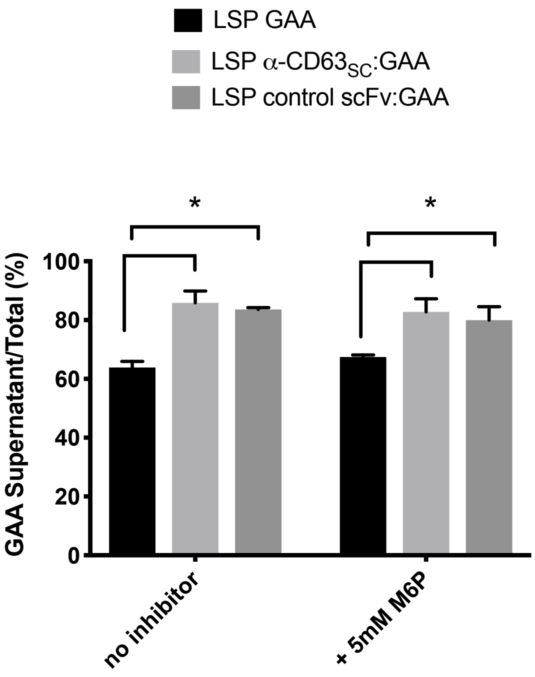

**Supplementary Figure S8**. Higher secreted to intracellular ratio of antibody:GAA versus GAA alone in Huh-7 hepatocytes. Huh-7 human hepatocytes were transiently transfected with liver-specific promoter (LSP) driven constructs encoding for GAA (black bars), α-CD63$_{SC}$:GAA (light gray bars), or a non-binding scFv:GAA control fusion protein (dark gray bars). Both scFv:GAA constructs had a higher ratio of protein in the secreted supernatant than GAA alone 3 days after transfection. Addition of M6P into the supernatant during the experimental period to mitigate CI-MPR-mediated uptake did not affect the ratio. * = $p < 0.05$, n=3

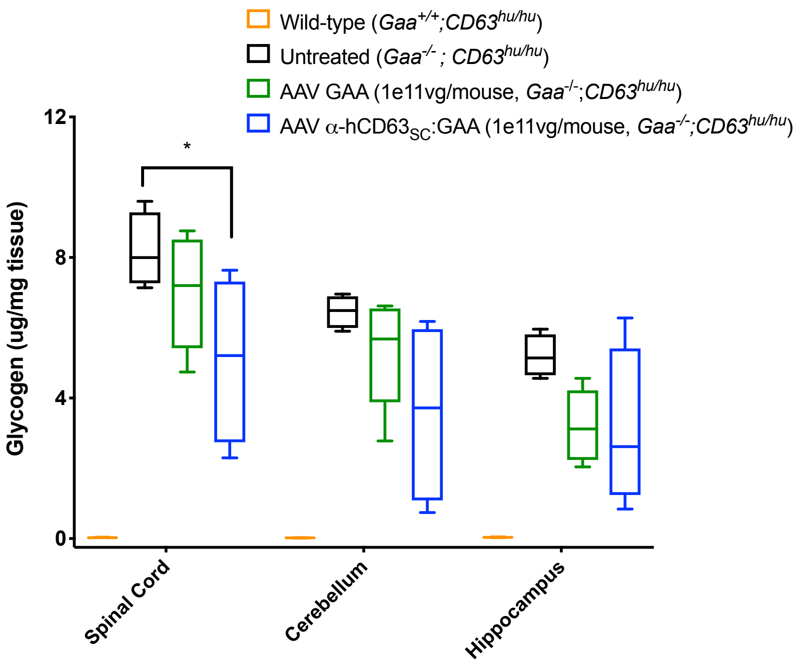

**Supplementary Figure S9.** Modest glycogen reductions in brain 10 months after treatment by AAV liver depot gene therapy. Reductions in glycogen were seen in both GAA and α-$CD63_{SC}$:GAA treated groups, but not in a statistically significant manner except for the spinal cord. * = $p < 0.05$

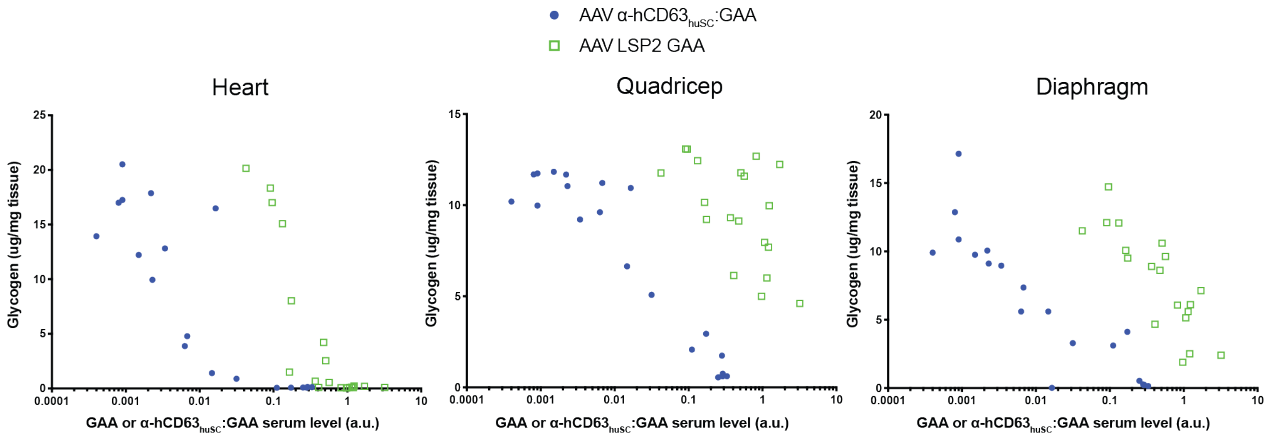

**Supplementary Figure S10.** α-hCD63$_{huSC}$:GAA clears more glycogen than GAA at equivalent serum levels. PD mice were treated with AAV α-hCD63$_{huSC}$:GAA or AAV-LSP2 GAA or at doses ranging from 5e11vg/kg to 4e12vg/kg to plot glycogen reduction by serum level. After 2 month of treatment, glycogen levels in heart, quadriceps, and diaphragm tissue lysates were plotted against the logarithm of the corresponding western blot serum level of the constructs.

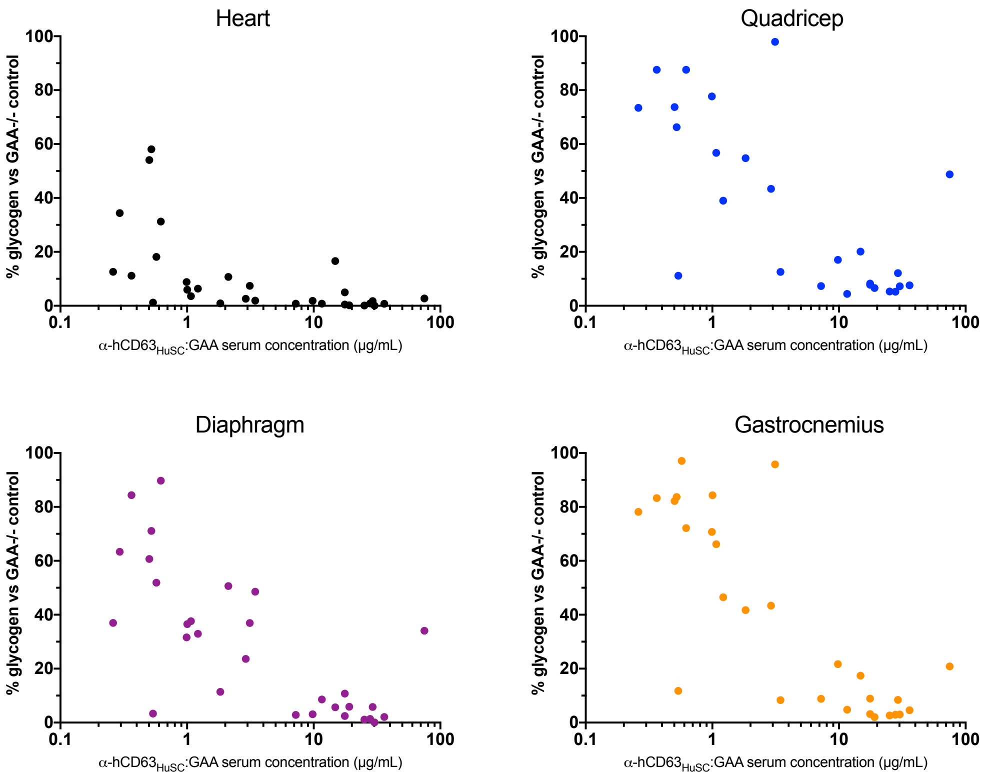

**Supplementary Figure S11.** Wild-type levels of glycogen are achieved in skeletal muscle at concentrations greater than ~5μg/mL of α-hCD63$_{huSC}$:GAA in the serum after 1 month of AAV treatment. Comparison of glycogen levels to ELISA-based quantification of serum levels of α-hCD63$_{huSC}$:GAA after 1 month of AAV treatment.

**Supplementary Methods**

*Generation of a $Cd63^{lacZ/lacZ}$ reporter mouse*
To generate allele VG13419 ($Cd63^{+/lacZ}$), a strategy similar to that used to create VG4323 ($Gaa^{-/-}$) was employed. The entire *Cd63* open reading frame, starting from the ATG in exon 3 (ENSMUSE00000150024) (Mouse genome coordinates 128910406-128912841, Chromosome 10), as well as 101bp of the 3' untranslated region was deleted and replaced by a lacZ insertion such that lacZ was in frame with the *Cd63* start ATG. Mouse BAC BMQ-373D17 was modified for use, creating 50 and 19 Kb homology arms, respectively. A fLoxed neomycin selection cassette was inserted following the lacZ. A GenBank file with the sequence of the targeting construct is supplied as supplementary materials. This targeting vector was then linearized with *Not*I and electroporated into 100% C57Bl/6NTac ESC. Cell clones positive for *Cd63* deletion were then electroporated with a Cre recombinase-containing plasmid to remove the resistance cassette. Following microinjection, $F_0$ offspring were bred to homozygosity ($Cd63^{lacZ/lacZ}$) on a 100% C57Bl/6NTac background for further study.

*Plasmid constructs of AAV-LSP2 GAA and AAV* α-$hCD63_{huSC}$*:GAA*
Fully human anti-human CD63 antibodies that bound to the 2nd extracellular loop of human CD63 were developed using Velocigene™ technologies. The variable domains were reformatted into an scFv and replaced the scFv portion in AAV α-$hCD63_{huSC}$:GAA using isothermal assembly. LSP2 GAA, a modified GAA with improved expression and secretion, was obtained from patent WO2018046774 (Sequence ID number 25) and cloned into an AAV expression plasmid.

*Hydrodynamic delivery of plasmid constructs into GAA knockout mice*
GAA and antibody:GAA plasmids were delivered into mice by intravenous hydrodynamic delivery. The GAA expression plasmid contained the human ubiquitin C promoter (NG_027722.2) with a rabbit beta-globin intron (AH001222.2) followed by a Kozak sequence and the cDNA from human GAA (NM_000152.4). Antibody:GAA plasmids consisted of a heavy chain:GAA fusion and a light chain plasmid. The heavy chain plasmid contained the human ubiquitin C promoter (NG_027722.2) with a rabbit beta-globin intron (AH001222.2) followed by a Kozak sequence, and codon modified cDNA for mROR1 signal peptide (BAA75480.1), the variable heavy domain of the antibody, the Fc of human IgG4 (P01861.1) with an S228P mutation, a GGGGS linker, and amino acids 70-952 of human GAA (NM_000152.4). The light chain plasmid consisted of ubiquitin C promoter (NG_027722.2) with a rabbit beta-globin intron (AH001222.2) followed by a Kozak sequence, and codon modified cDNA for mROR1 signal peptide (BAA75480.1), the variable light domain of the antibody, and the human constant kappa light chain (P01834.2).To express GAA, 40μg of the GAA plasmid was injected. To express antibody:GAA, 40μg of antibody:GAA and 40μg of the corresponding light chain were injected. Serum was collected on a regular basis by tail-nick bleeds. Mice were sacrificed 3 weeks post-HDD.

*Secretion of* α-$hCD63_{SC}$*:GAA and GAA in Huh-7 hepatocytes*

Huh-7 cells were transfected with LSP GAA (n=3), LSP α-hCD63$_{huSC}$:GAA(n=3), or a non-binding LSP control scFv:GAA(n=3) plasmid using Mirus TransIT-LT1 transfection reagent per manufacturer's instructions. This experiment was repeated with competitive inhibitors against for CI-MPR and hCD63 by performing the experiment in the presence of with 5mM mannose 6-phosphate and 200nM anti-hCD63 antibody. Cells were lysed 3 days after transfection with RIPA buffer and analyzed by western blot for GAA and hGAPDH (Abcam, ab9484). Total secretion was calculated as a ratio of total GAA signal in the supernatant to total GAA signal in the supernatant and lysate.

*Purification of GAA*
GAA was expressed in CHO-K1 cells by transiently transfecting a plasmid encoding an mROR signal peptide with a myc tag (EQKLISEEDL) followed by amino acids 70-952 of human GAA. Supernatants were collected 72 hours post transfection and bound on a myc column (ThermoFisher #20168) and eluted using myc peptide (ThermoFisher #20170).

*Glycan analysis of antibody:GAA fusions*
The N-linked glycans on α-hCD63:GAA protein were released by treatment with PNGase F, followed by derivatization of the reduced glycan chain termini with the fluorescent reagent, 2-Anthranilic Acid (2-AA). The AA-labeled glycans were then separated with a Waters BEH glycan column and monitored using a fluorescence detector at an excitation wavelength of 360 nm and emission wavelength of 420 nm. A Thermo Q-Exactive mass spectrometer was used to collect the on-line MS and MS/MS data for glycan structure identification.

*scFv:GAA serum concentration ELISA assay*

A sandwich ELISA was developed to measure the concentration of α-hCD63$_{huSC}$:GAA in serum. Briefly, high protein-binding 96-well plates were coated with 2μg/mL of a monoclonal antibody that recognized the light chain of the scFv. Plates were blocked with bovine serum albumin. Dilutions of mouse serum or α-hCD63$_{huSC}$:GAA protein standards were incubated on the plate overnight. Bound α-hCD63$_{huSC}$:GAA was detected with recombinant protein encoding the 2$^{nd}$ extracellular loop domain of human CD63 (that the parental antibody to the anti-hCD63 antibody was raised against) tagged to human Fc. An HRP-goat anti-human Fc (Jackson Immunoresearch, 709-035-098) and colorimetric detection kit was used to quantify bound α-hCD63$_{huSC}$:GAA.
